# Phosphorylation of the selective autophagy receptor TAX1BP1 by canonical and noncanonical IκB kinases promotes its lysosomal localization and clearance of MAVS aggregates

**DOI:** 10.1101/2021.01.07.425702

**Authors:** Young Bong Choi, Jiawen Zhang, Mai Tram Vo, Jesse White, Chaoxia He, Edward W. Harhaj

## Abstract

TAX1BP1 is a selective autophagy receptor which inhibits NF-κB and RIG-I-like receptor (RLR) signaling to prevent excessive inflammation and maintain homeostasis. Selective autophagy receptors such as p62/SQSTM1 and OPTN are phosphorylated by the noncanonical IκB kinase TBK1 to stimulate their selective autophagy function. However, it is unknown if TAX1BP1 is regulated by TBK1 or other kinases under basal conditions or during RNA virus infection. Here, we found that the noncanonical IκB kinases TBK1 and IKKi phosphorylate TAX1BP1 to regulate its basal turnover, whereas the canonical IκB kinase IKKα and the core autophagy factor ATG9 play essential roles in RNA virus-mediated TAX1BP1 autophagosomal degradation. TAX1BP1 phosphorylation by canonical and noncanonical IκB kinases promotes its localization to lysosomes resulting in its degradation. Furthermore, TAX1BP1 plays a critical role in the clearance of MAVS aggregates, and phosphorylation of TAX1BP1 augments its MAVS aggrephagy function. Together, our data support a model whereby IκB kinases license TAX1BP1 selective autophagy function to inhibit MAVS and RLR signaling.

**Author Summary:** The RIG-I-like receptor (RLR) pathway induces type I interferon (IFN) and proinflammatory cytokines in response to RNA virus infection. MAVS is a mitochondrial adaptor protein in the RLR pathway that forms prion-like aggregates upon activation; however, how MAVS aggregates are cleared to restore homeostasis is unclear. Autophagy is a lysosomal degradation pathway important for the clearance of potentially cytotoxic protein aggregates that could induce inflammation and/or cell death. TAX1BP1 is a selective autophagy receptor that inhibits RLR signaling, but the precise mechanisms remain unknown. Here, we found that TAX1BP1 is a substrate for multi-site phosphorylation by canonical and noncanonical IκB kinases which triggered its lysosomal localization and degradation. We also found that TAX1BP1 was critical for the clearance of MAVS aggregates in a phosphorylation-dependent manner. Overall, our data suggest that phosphorylation serves a key regulatory function for TAX1BP1 to inhibit RLR signaling.

## Introduction

Pattern recognition receptors (PRRs) detect conserved molecular features of viruses and other pathogens known as PAMPs (pathogen-associated molecular patterns). In the RIG-I-like receptor (RLR) pathway the cytoplasmic RNA helicases RIG-I and MDA5 recognize nucleic acid derived from RNA viruses and trigger signaling pathways through the mitochondrial protein MAVS, leading to activation of NF-κB and IRF3 transcription factors that upregulate expression of proinflammatory cytokines and type I IFNs respectively [1, 2]. MAVS recruits E3 ubiquitin ligases such as TRAF2, TRAF3, TRAF5 and TRAF6 to conjugate lysine 63 (K63)-linked polyubiquitin chains that recruit the adaptor NEMO (also known as IKKγ) and noncanonical IκB kinases TBK1 and IKKi (also known as IKK epsilon) [3–8]. TBK1 and IKKi directly phosphorylate IRF3 and IRF7 to trigger their dimerization, nuclear localization and activation of type I IFN to restrict virus replication [9, 10]. In addition, canonical IKK kinases IKKα and IKKβ phosphorylate the inhibitory protein IκBα to trigger its proteasomal degradation and release NF-κB dimers to activate proinflammatory genes [11].

Macroautophagy (hereafter referred to as autophagy) is an evolutionarily conserved lysosomal degradation pathway critical for homeostasis. Autophagy is initiated by the recruitment of membranes and the formation of a phagophore which elongates and forms double membrane vesicles termed autophagosomes which subsequently fuse with lysosomes to degrade the contents in autolysosomes [12, 13]. LC3-I is an ATG8 family member conjugated with phosphatidylethanolamine (PE) during autophagy to form LC3-II and plays key roles in the biogenesis and maturation of autophagosomes as well as cargo recruitment [14]. Selective autophagy results in the recruitment of specific cargo to autophagosomes including protein aggregates/misfolded proteins (aggrephagy), damaged organelles such as mitochondria (mitophagy) or pathogenic microbes (xenophagy) [15]. Autophagy receptors provide specificity by recognizing and linking cargo to autophagosomes. Cargo destined for autophagosomes are typically modified by post-translational modifications (PTMs) such as ubiquitination which can be detected by autophagy receptors containing ubiquitin binding domains [16]. Furthermore, autophagy receptors harbor LC3 interaction regions (LIRs) that link cargo to autophagosomes [17]. The best characterized selective autophagy receptors consist of the Sequestosome 1 (p62-SQSTM1)-like receptor family including p62, Optineurin (OPTN), NBR1, NDP52 and TAX1BP1 [18]. Autophagy has been linked to the negative regulation of RLR signaling [19, 20], yet the precise mechanisms remain unknown.

TAX1BP1 was originally identified in yeast two-hybrid screens as a binding protein of the human T-cell leukemia virus 1 (HTLV-1) Tax protein, the ubiquitin-editing enzyme A20 (also known as TNFAIP3) and the E3 ubiquitin ligase TRAF6 [21–23]. TAX1BP1 inhibits canonical NF-κB signaling, together with E3 ligases Itch and RNF11, by acting as an adaptor for the ubiquitin-editing enzyme A20 [24–27]. In addition to regulating NF-κB signaling, TAX1BP1 also inhibits the RLR pathway and the induction of type I IFN triggered by RNA virus infection or transfection with the double-stranded RNA mimetic poly(I:C) [28]. Furthermore, TAX1BP1 blocks RLR-mediated apoptosis by interacting with and promoting MAVS degradation [29]. TAX1BP1 also suppresses the TLR3/4 pathways by targeting the adaptor TRIF for degradation [30, 31]. TAX1BP1 contains two LIR motifs and functions as a selective autophagy receptor [32–34]. Furthermore, the second zinc finger domain (ZnF2) in the carboxyl-terminus of TAX1BP1 can bind to K63-linked polyubiquitin chains [34, 35]. Therefore, TAX1BP1 targets ubiquitinated cargo via ZnF2 and recruits cargo to developing autophagosomes via the LIR domains. TAX1BP1 also interacts with myosin VI, a cytoskeletal actin-based motor protein regulating vesicular transport, to induce autophagosome maturation [34]. Therefore, TAX1BP1 exerts multiple roles in autophagy including cargo selection and autophagosome maturation. TAX1BP1 can remove damaged mitochondria (mitophagy) together with OPTN and NDP52 [36], and pathogenic bacteria including *Salmonella Typhimurium* and *Mycobacterium tuberculosis* (xenophagy) [34, 37]. A recent study has linked TAX1BP1 to the clearance of protein aggregates (i.e., polyQ huntington fragments and TDP-43) in the brain thus implicating TAX1BP1 as an aggrephagy receptor [38].

Despite the important roles of TAX1BP1 in the inhibition of innate immune signaling pathways, it remains unclear how the selective autophagy function of TAX1BP1 is regulated and if TAX1BP1 functions as an aggrephagy receptor in the regulation of innate immunity. We previously reported that phosphorylation of TAX1BP1 by the kinase IKKα plays a critical role in the termination of TNF and IL-1β-induced NF-κB signaling [39]; however, it is unknown if phosphorylation of TAX1BP1 regulates its autophagy function. In this study, we have identified the noncanonical IκB kinases TBK1 and IKKi as regulators of TAX1BP1 basal autophagic degradation. However, during RNA virus infection, both TBK1 and IKKi are dispensable for TAX1BP1 degradation, whereas IKKα and the core autophagy factor ATG9 play critical roles in the inducible autophagic degradation of TAX1BP1. Furthermore, TAX1BP1 mediates the clearance of MAVS aggregates, both basally and during RNA virus infection, and phosphorylation of TAX1BP1 stimulates its MAVS aggrephagy function.

## Results

### TAX1BP1 is phosphorylated by IKKi and TBK1 kinases

We previously reported that the IKKα subunit of IKK phosphorylates TAX1BP1 to promote the termination of NF-κB signaling [39]. During the course of our studies on TAX1BP1 regulation of RLR signaling, we unexpectedly found that the noncanonical IκB kinases TBK1 and IKKi also phosphorylated TAX1BP1. Overexpression of IKKα, IKKi and TBK1 all induced a slower migrating form of TAX1BP1 (Fig 1A). Interestingly, IKKi (and to a lesser extent TBK1) overexpression was associated with the loss of TAX1BP1 protein (Fig 1A), likely due to its degradation. Treatment with lambda phosphatase converted the slower migrating form to a faster migrating form of TAX1BP1, thus confirming phosphorylation (Fig 1B). Furthermore, a kinase dead IKKi mutant K38A was impaired in TAX1BP1 phosphorylation (Fig S1). *In vitro* kinase assays with purified recombinant proteins demonstrated that TBK1 and IKKi could both directly phosphorylate TAX1BP1, although IKKi induced a more obvious TAX1BP1 band shift (Fig 1C). Bioinformatics analysis using NetPhos2.0 revealed two putative IKKi phosphorylation sites at Ser254 and Ser593 in TAX1BP1, similar to a site found in the deubiquitinase CYLD (Fig 1D) [40]. Interestingly, these two sites were also identified in our previous study on IKKα-induced TAX1BP1 phosphorylation in the context of NF-κB signaling [39]. To identify the IKKi-inducible TAX1BP1 phosphorylation sites in an unbiased manner, we used liquid chromatography coupled with tandem mass spectrometry (LC-MS/MS). As expected, IKKi induced a slower migrating band shift in TAX1BP1 (Fig 1E). A total of four bands were excised from the gel (both phosphorylated and unphosphorylated TAX1BP1 as controls) and subjected to LC-MS/MS analysis, which identified a total of 13 IKKi-inducible TAX1BP1 phosphorylation sites in TAX1BP1 (Fig 1F).

**Fig. 1.**
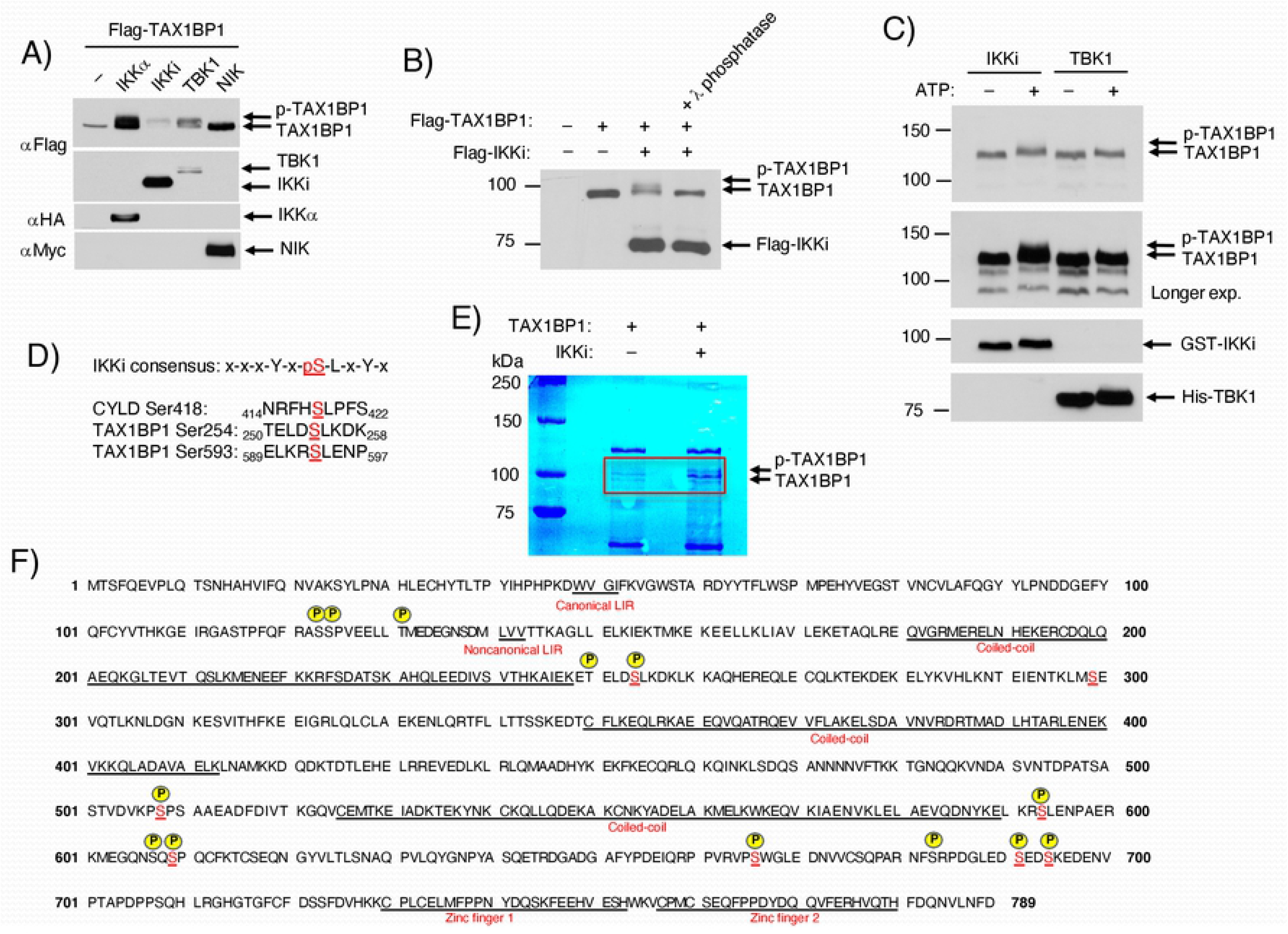
IκB kinases induce TAX1BP1 phosphorylation. (A) 293T cells were co-transfected with Flag-TAX1BP1 together with the indicated kinase plasmids and 24 h later lysed, and the cell extracts were immunoblotted with the indicated antibodies. (B) 293T cells were transfected with the indicated plasmids and lysates were incubated with λ-phosphatase for 30 min prior to immunoblotting with anti-FLAG. (C) *In vitro* kinase assays. Purified GST-tagged TAX1BP1 (300 ng) was incubated with 50 ng of purified recombinant GST-tagged IKKi or hexahistidine-tagged TBK1 in the presence or absence of ATP. The reaction mixtures were immunoblotted with antibodies to TAX1BP1, IKKi and TBK1. (D) Consensus phosphorylation sequences for IKKi found in CYLD and predicted for TAX1BP1. The conserved serine residue is highlighted in red. (E) Colloidal blue staining of *in vitro* kinase reaction mixtures containing TAX1BP1 with or without IKKi. Individual bands within the red rectangle were cut and gel extracted for mass spectrometry (MS) analysis. (F) The primary sequences of TAX1BP1. Thirteen serine and threonine residues were predicted for IKKi phosphorylation of TAX1BP1 by MS analysis and are marked by yellow circles above them. The amino acids for the main functional domains of TAX1BP1, including canonical and noncanonical LC3-interacting regions (LIRs), three coiled-coil domains, and two zinc-finger domains, are underlined.

### Mapping of TAX1BP1 phosphorylation sites

To validate the putative TAX1BP1 phosphorylation sites, we generated a panel of TAX1BP1 point mutants with all 13 putative sites mutated to alanine (designated as 13A) as well as the 10 sites (designated as 10A) downstream of coiled coil domain 1 (CC1) (Fig 2A). IKKi overexpression induced the phosphorylation of WT TAX1BP1, but not of 13A or 10A mutants (Fig 2B). Since the TAX1BP1 10A mutant was indistinguishable from 13A with regard to the lack of phosphorylation and degradation we focused on this mutant for subsequent experiments. We next generated a new panel of rescue mutants where each of the potential phosphorylation sites was individually restored back to the original amino acid in the context of TAX1BP1 10A (Fig 2C). These are designated as TAX1BP1 9A/WT amino acid. This panel of TAX1BP1 mutants was transfected into cells and then infected with an RNA virus, VSV-GFP (vesicular stomatitis virus (VSV) encoding a green fluorescence protein (GFP) reporter), followed by western blotting to assess phosphorylation. As expected, WT TAX1BP1 was phosphorylated and degraded upon VSV infection (Fig 2D). However, TAX1BP1 10A was resistant to VSV-induced phosphorylation and degradation (Fig 2D). VSV-induced phosphorylation was observed with TAX1BP1 9A/T250, 9A/S254 and 9A/S593, of which S254 and S593 are predicted IKKi phosphorylation sites. We also observed IKKi-induced phosphorylation of TAX1BP1 9A/S666 (Fig S2). Since virus-induced TAX1BP1 degradation was not fully restored with single point mutations, we next generated a TAX1BP1 compound mutant with S254, S593 and S666 in the context of 10A (designated as 7A; Fig 2C). An additional mutant was generated with T250, S254, S593 and S666 in the context of 10A (designated as 6A; Fig 2C). Remarkably, VSV-induced TAX1BP1 degradation was restored by the 7A and 6A mutants (Fig 2E) suggesting that S254, S593 and S666 (and possibly T250) act redundantly in promoting TAX1BP1 phosphorylation/degradation. We also introduced point mutations in both canonical (W49A) and noncanonical (V143S) LC3 interaction motifs in TAX1BP1. Although IKKi-induced phosphorylation was not affected by the single and double TAX1BP1 LC3 binding mutants, the degradation of these mutants was impaired (Fig 2F). Thus, both TAX1BP1 phosphorylation and LC3 interaction motifs regulate its degradation.

**Fig. 2.**
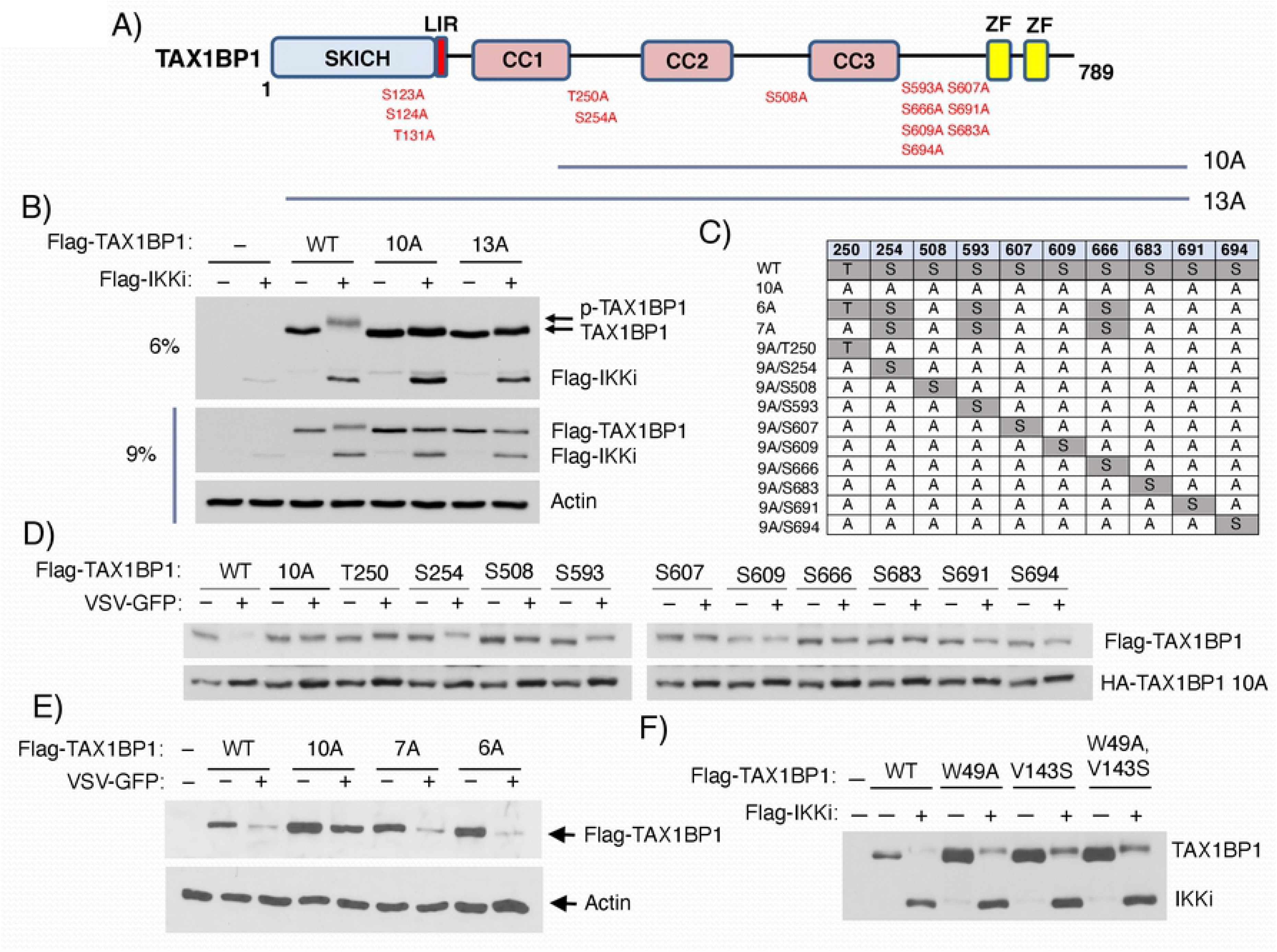
Three phosphoserine residues are involved in TAX1BP1 degradation. (A) Schematic diagram of TAX1BP1 domains. The marked ten and thirteen predicted phosphorylation sites were substituted for alanine, generating the TAX1BP1 10A and 13A mutants, respectively. SKICH, the SKIP carboxy homology domain; LIR, LC3-interacting region; CC, coiled-coil domain; ZF, zinc finger domain. (B) Immunoblot analysis of the extracts derived from 293T cells transfected with Flag-TAX1BP1 wild type (WT) and mutants (10A and 13A) together with or without Flag-IKKi. For better separation of phosphorylated TAX1BP1, a 6% gel was used. (C) Schematic of restored TAX1BP1 10A variants at single or multiple phosphorylation residue(s). (D) 293T cells transfected with Flag-TAX1BP1 WT and mutants together with HA-tagged TAX1BP1 10A for 24 h were infected with or without VSV-GFP for 6 h at an MOI of 1, and the cell extracts were immunoblotted with the indicated antibodies. (E) Immunoblot analysis of the extracts derived from 293T cells transfected with Flag-TAX1BP1 WT and variants (10A, 7A, and 6A) and 24 h later infected with or without VSV-GFP as above. As shown in (C), 7A was generated by restoring the three potential phosphorylation sites, A254, A593, and A666 of 10A, and 6A was generated by restoring A250 of 7A. (F) 293T cells were co-transfected with Flag-TAX1BP1 WT and LIR mutants, W49A for mutation of the canonical LIR, V143S for mutation of the non-canonical LIR, and W49A/V143S for mutation of both the LIRs, together with or without Flag-IKKi for 24 h, and the cell extracts were immunoblotted with the indicated antibodies.

### TBK1 and IKKi regulate the basal phosphorylation and turnover of TAX1BP1

We hypothesized that virus infection-mediated phosphorylation and autophagic degradation of TAX1BP1 was mediated by IKKi. To test this notion, we generated IKKi knockout (KO) DLD-1 cell lines using CRISPR/Cas9 technology. In addition, we generated DLD-1 cell lines deficient in the closely related kinase TBK1 due to their functional redundancy. DLD-1 cells were used because of high basal levels of TAX1BP1 expression [29]. DLD-1 cells were transduced with recombinant lentiviruses expressing Cas9 and either IKKi or TBK1 gRNAs followed by limiting dilution and clonal analysis of TBK1 and IKKi knockouts. Multiple clones of TBK1 and IKKi knockouts were identified and two clones each were selected for further experimentation. Surprisingly, poly(I:C)-induced TAX1BP1 degradation remained intact in TBK1 or IKKi KO cells (Fig 3A and 3B). To address the possibility of functional compensation between TBK1 and IKKi, we also generated TBK1/IKKi double knockout (TBK1/IKKi dKO) DLD-1 cells and infected these cells with VSV-GFP. Similarly, TAX1BP1 degradation was unimpaired in TBK1/IKKi dKO cells infected with VSV-GFP (Figs 3C and S3). Interestingly, VSV-induced p62/SQSTM1, but not NDP52, degradation was partially impaired in TBK1/IKKi dKO cells (Fig 3C). TAX1BP1 degradation also remained intact in VSV-GFP-infected *Ikki*^−/−^ and *Ikki^−/−^Tbk1^−/−^* MEFs (Fig 3D), thus ruling out cell-type specific effects. During the course of these studies, we noticed that basal expression of TAX1BP1 protein was increased in TBK1/IKKi dKO cells which was confirmed by quantification in several independent experiments (Fig 3E). To provide further evidence that the basal phosphorylation of TAX1BP1 caused its degradation, we treated cells with the phosphatase inhibitor calyculin A. Indeed, calyculin A promoted TAX1BP1 degradation in WT DLD-1 cells, which was partially impaired in TBK1/IKKi dKO cells (Fig 3F). Therefore, it appears that dynamic regulation of TAX1BP1 phosphorylation and dephosphorylation controls its turnover.

**Fig. 3.**
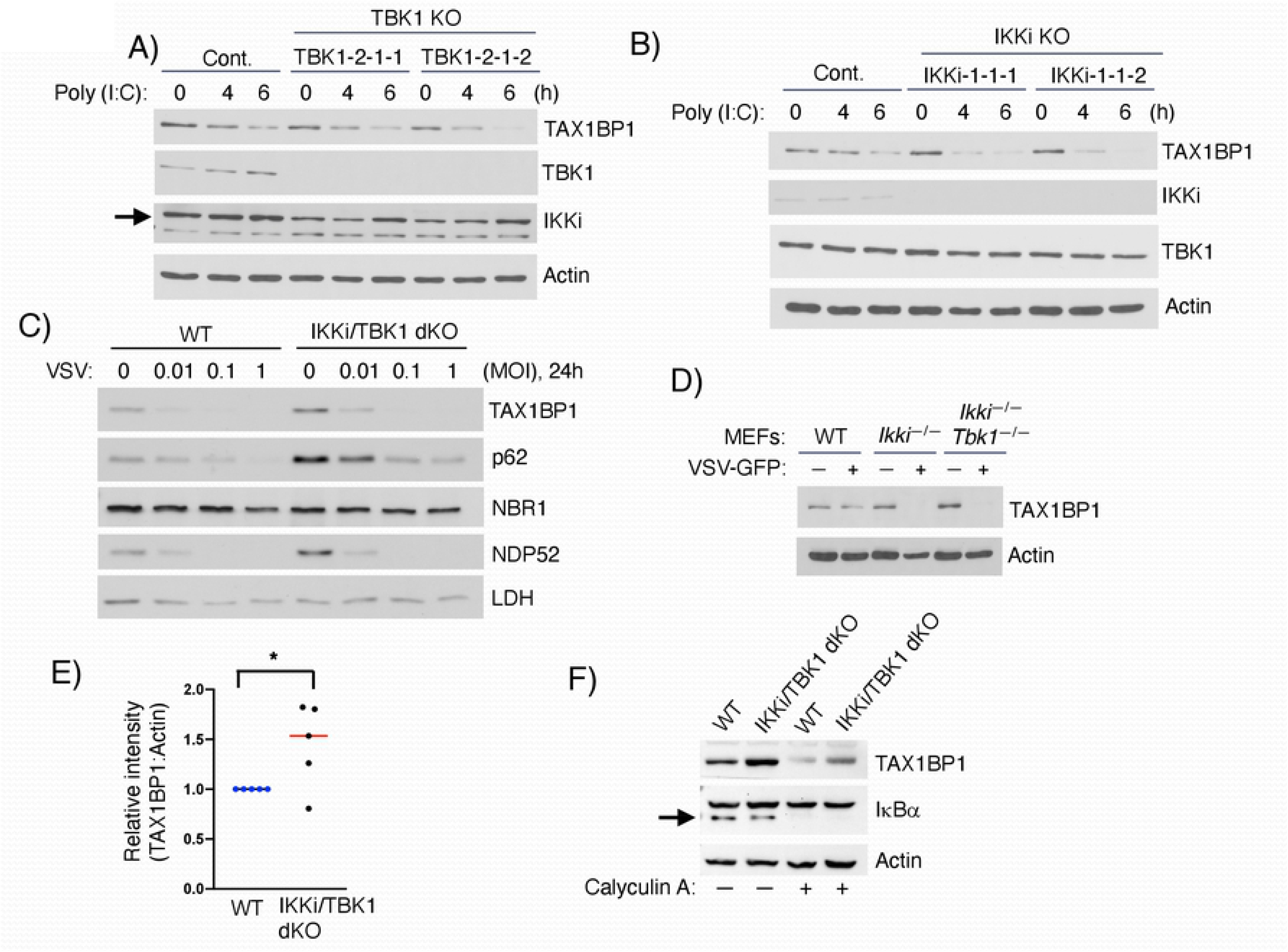
TBK1 and IKKi regulate the basal turnover of TAX1BP1. (A-D) Immunoblot analyses of the extracts derived from the following cells: two different *TBK1* knockout (KO) DLD-1 cell lines (A) and two different *IKKi* KO DLD-1 cell lines (B) transfected with 2.5 μg/ml poly(I:C) for 0, 4 and 6 h, *IKKi/TBK1* double KO (dKO) DLD-1 cells infected with VSV-GFP for 24 h at different MOIs (C), and *Ikki*^−/−^ and *Ikki^−/−^ Tbk1^−/−^* MEFs infected with VSV-GFP for 20 h at an MOI of 0.1 (D). (E) TAX1BP1 expression was quantified by ImageJ using lysates from WT and *IKKi/TBK1* dKO DLD-1 cells. Data were derived from five independent experiments. Unpaired Student’s *t*-test, \**P* <0.05. (F) WT and *IKKi/TBK1* dKO DLD-1 cells were treated with calyculin A for 30 min, and lysates were immunoblotted with the indicated antibodies.

### IKKα is required for virus-triggered degradation of TAX1BP1

We previously reported that IKKα can directly phosphorylate TAX1BP1 to promote its NF-κB inhibitory function [39]. However, it remains unclear if IKKα plays a role in virus infection-induced autophagic degradation of TAX1BP1. Therefore, we first pretreated control and IKKi/TBK1 dKO DLD-1 cells with a small molecule IKK inhibitor (IKK inhibitor VII) which inhibits both IKKα (IC_50_=40 nM) and IKKβ (IC_50_=200 nM). Poly(I:C) induced the degradation of TAX1BP1 in vehicle-treated WT and IKKi/TBK1 dKO DLD-1 cells, but not in WT and IKKi/TBK1 dKO DLD-1 cells treated with IKK inhibitor VII (Fig 4A). To determine which IKK subunit was required for virus-induced TAX1BP1 degradation, WT, *Ikkα^−/−^, Ikkβ*^−/−^ and *Ikkγ*^−/−^ MEFs were infected with VSV-GFP and TAX1BP1 expression was examined by western blotting. Interestingly, VSV-induced TAX1BP1 degradation was only impaired in *Ikkα*^−/−^ MEFs (Fig 4B). Together, these data suggest that IKKα is required for virus-induced degradation of TAX1BP1.

**Fig. 4.**
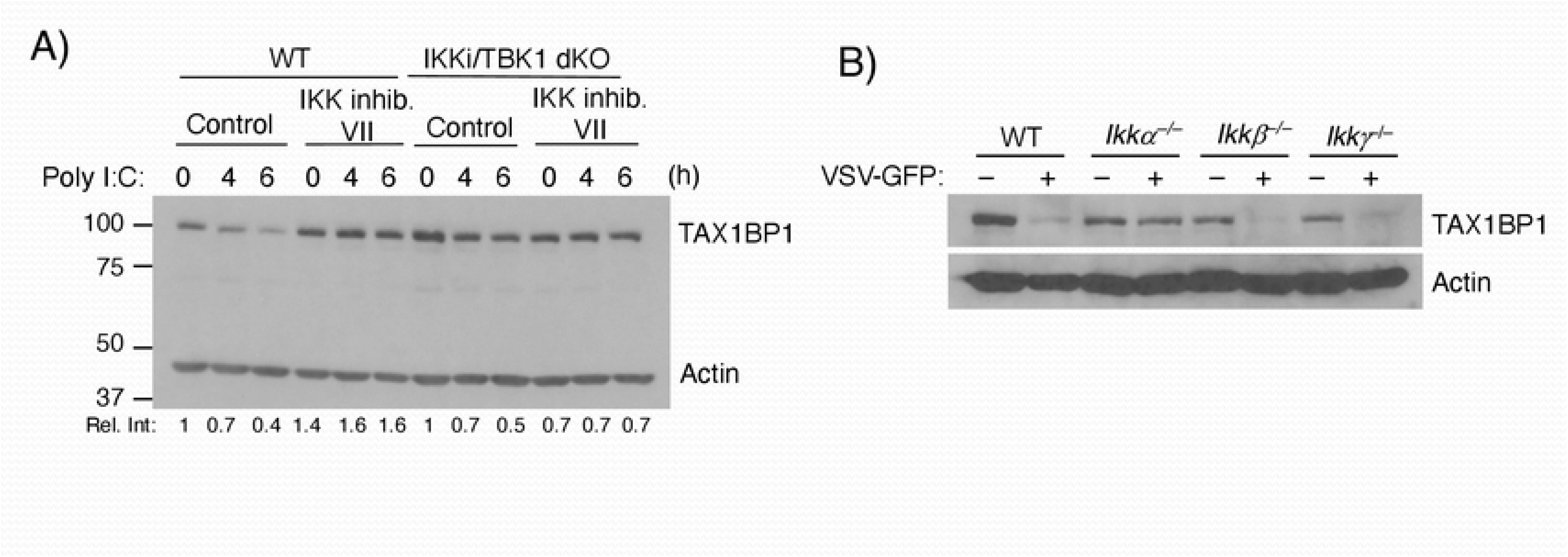
IKKα is required for VSV-induced TAX1BP1 degradation. (A) WT and *IKKi/TBK1* dKO DLD-1 cells were treated with DMSO or 20 μM IKK inhibitor VII for 1 h before poly(I:C) transfection. Lysates were immunoblotted with the indicated antibodies. (B) *Ikkα*^−/−^, *Ikkβ*^−/−^ and *Ikk*^−/−^ MEFs were infected with VSV-GFP for 13 h and TAX1BP1 and Actin expression were examined by immunoblotting.

### TAX1BP1 is degraded via autophagy during RNA virus infection

In the latter stages of autophagy, selective autophagy receptors are degraded in autolysosomes together with their cargo, and are often used as markers for autophagic flux. Therefore, TAX1BP1 phosphorylation during virus infection likely stimulates its autophagy function, which then triggers its own degradation in autolysosomes. To confirm that TAX1BP1 degradation elicited by RNA virus infection was mediated by autophagy we first treated DLD-1 cells with either vehicle or bafilomycin A1 (Baf A1), a specific inhibitor of vacuolar-type H+ ATPase (V-ATPase) that prevents the maturation of autophagic vacuoles. Although poly(I:C) triggered TAX1BP1 degradation as expected, Baf A1 blocked poly(I:C)-induced TAX1BP1 degradation (Fig 5A). ATG3 functions as an E2-like enzyme in the conjugation of PE to LC3-I to yield LC3-II. We examined TAX1BP1 degradation in WT and *Atg3*^−/−^ MEFs infected with VSV-GFP at a range of MOIs. TAX1BP1 degradation was induced by VSV-GFP in WT MEFs, which was largely impaired in *Atg3*^−/−^ MEFs, however there was still an appreciable amount of TAX1BP1 degradation suggesting potential degradation routes independent of LC3 lipidation (Fig 5B). As expected, virus infection induced the conversion of LC3-I to LC3-II in WT MEFs, but not in *Atg3* MEFs (Fig 5B). However, VSV-GFP infection in *Atg3*^−/−^ MEFs was comparable to WT MEFs as examined by Incucyte S3 live-cell analysis (Fig 5C). To further investigate the requirement of other autophagy components we used CRISPR/Cas9 to generate ATG9 and NCOA4 KO DLD-1 cell lines. ATG9 is a transmembrane protein required for autophagy that delivers membranes to expanding phagophores [41]. NCOA4 is a selective autophagy receptor and TAX1BP1 interacting protein that mediates the lysosomal degradation of ferritin to regulate iron homeostasis [42]. Degradation of TAX1BP1 was impaired in response to poly(I:C) transfection or VSV infection in clonal ATG9 KO cells (Fig 5D and 5E). However, NCOA4 deficiency had no effect on TAX1BP1 degradation (Fig 5D). Taken together, these data suggest that TAX1BP1 degradation induced by RNA virus infection is mainly mediated by a classical autophagy pathway dependent on ATG9 and LC3 lipidation.

**Fig. 5.**
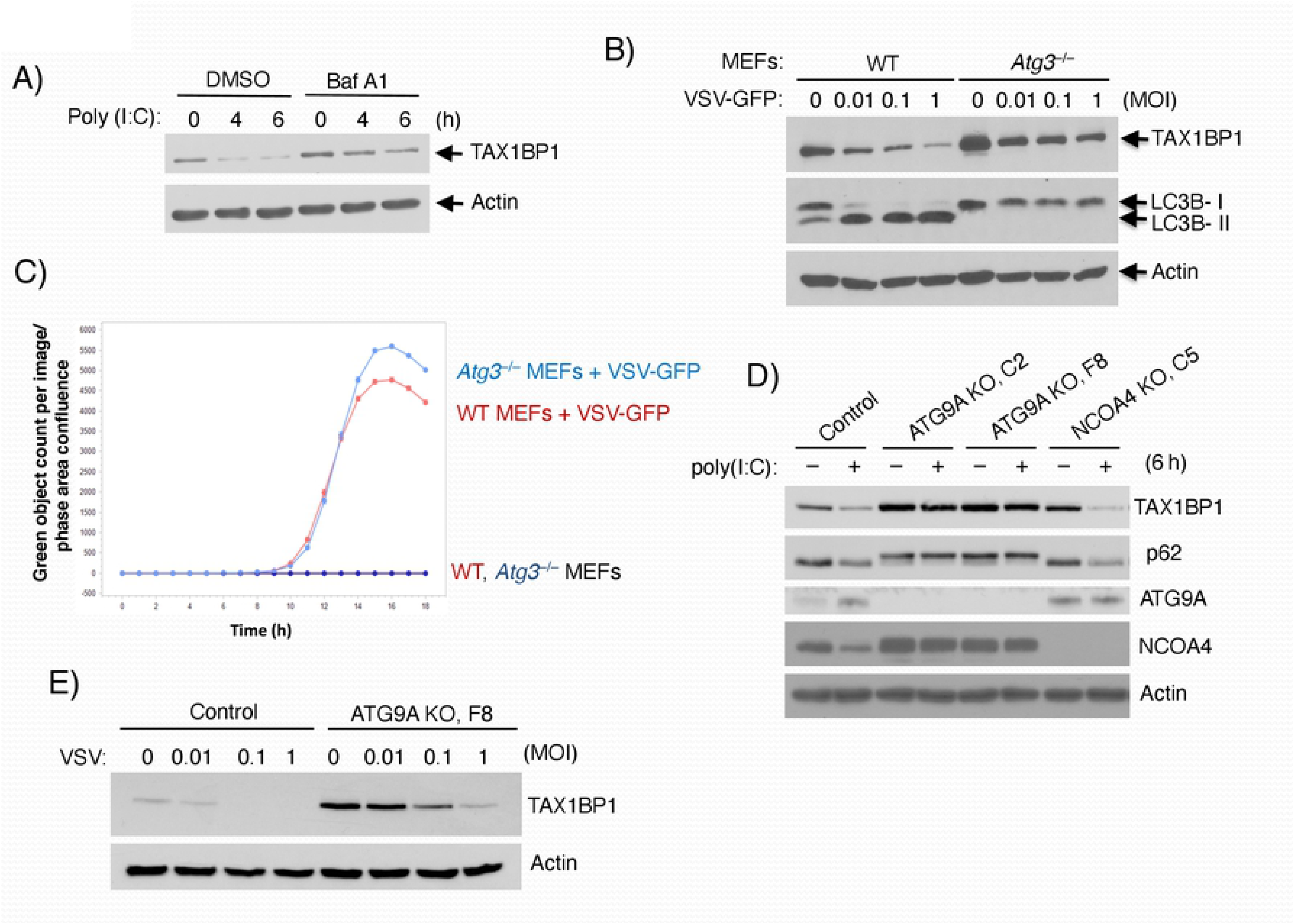
TAX1BP1 is degraded by autophagy during RNA virus infection. (A) DLD-1 cells were treated with DMSO or Baf A1 prior to poly(I:C) transfection. Lysates were subjected to immunoblotting with the indicated antibodies. (B) WT and *Atg*3^−/−^ MEFs were infected with VSV-GFP at the indicated MOIs for 13 h. (C) Incucyte S3 live-cell analysis was performed with WT and *Atg*3^−/−^ MEFs infected with VSV-GFP (MOI=0.1). (D) Immunoblot analyses of the extracts derived from two different *ATG9A* KO DLD-1 cell lines (C2 and F8) and *NCOA5* KO DLD-1 cell line (C5) transfected with 2.5 μg/ml poly(I:C) for 6 h. To facilitate the detection of ATG9A protein, cell lysates were prepared by heating at 70°C for 10 min in 1x SDS sample buffer instead of boiling. (E) *ATG9A* KO DLD-1 cell line (F8) was infected with VSV-GFP for 24 h at different MOIs and subjected to immunoblotting with the indicated antibodies.

### TAX1BP1 phosphorylation promotes its lysosomal localization

To determine how phosphorylation enhances TAX1BP1 autophagy function and degradation in autophagosomes, we first examined binding with ATG8 family members. TAX1BP1 could strongly interact with LC3A, LC3B, LC3C, GABARAP and GEC1, but weakly with GATE16 (Fig S4). However, TAX1BP1 phosphorylation mutants (10A, 7A and 6A), as well as the S254D, S593D and S666D phosphomimetic 3SD (serines 254, 593 and 666 mutated to aspartic acid residues) all similarly interacted with LC3B, GEC1 and MAVS (Figs S5 and S6). To determine if TAX1BP1 phosphorylation regulated its dimerization, we performed NanoBiT assays (Promega) with WT TAX1BP1, phosphorylation mutant 10A and phosphomimetic 3SD. However, the 10A and 3SD mutants exhibited comparable dimerization to WT TAX1BP1 (Fig S7). Therefore, TAX1BP1 phosphorylation does not appear to regulate LC3 or MAVS binding, as well as its dimerization.

We next sought to determine if phosphorylation regulated the localization of TAX1BP1 to lysosomes. To this end, we examined the subcellular localization of WT TAX1BP1, the phosphorylation mutant 10A, and phosphomimetic 3SD in transfected *TAX1BP1* KO HeLa cells. Cells were treated with the protease inhibitor leupeptin to inhibit autophagic flux and prevent TAX1BP1 degradation by poly(I:C). WT TAX1BP1 colocalization with LAMP1, a marker of lysosomes, was significantly increased by poly(I:C) transfection (Fig 6A and 6B). However, the phosphorylation-deficient 10A mutant was impaired in poly(I:C)-mediated colocalization with lysosomes (Fig 6A and 6B). Remarkably, the TAX1BP1 phosphomimetic 3SD colocalized persistently with lysosomes, which was not further increased by poly(I:C) (Fig 6A and 6B). To determine the role of IκB kinases in TAX1BP1 lysosomal targeting, we overexpressed IKKα, IKKβ and IKKi with TAX1BP1 and stained for LAMP1. Overexpression of IKKα and IKKi, but not IKKβ, promoted TAX1BP1 colocalization with LAMP1 and lysosomes (Fig S8). Therefore, IKKα- and IKKi-mediated TAX1BP1 phosphorylation at serines 254, 593 and 666 directs its localization to lysosomes.

**Fig. 6.**
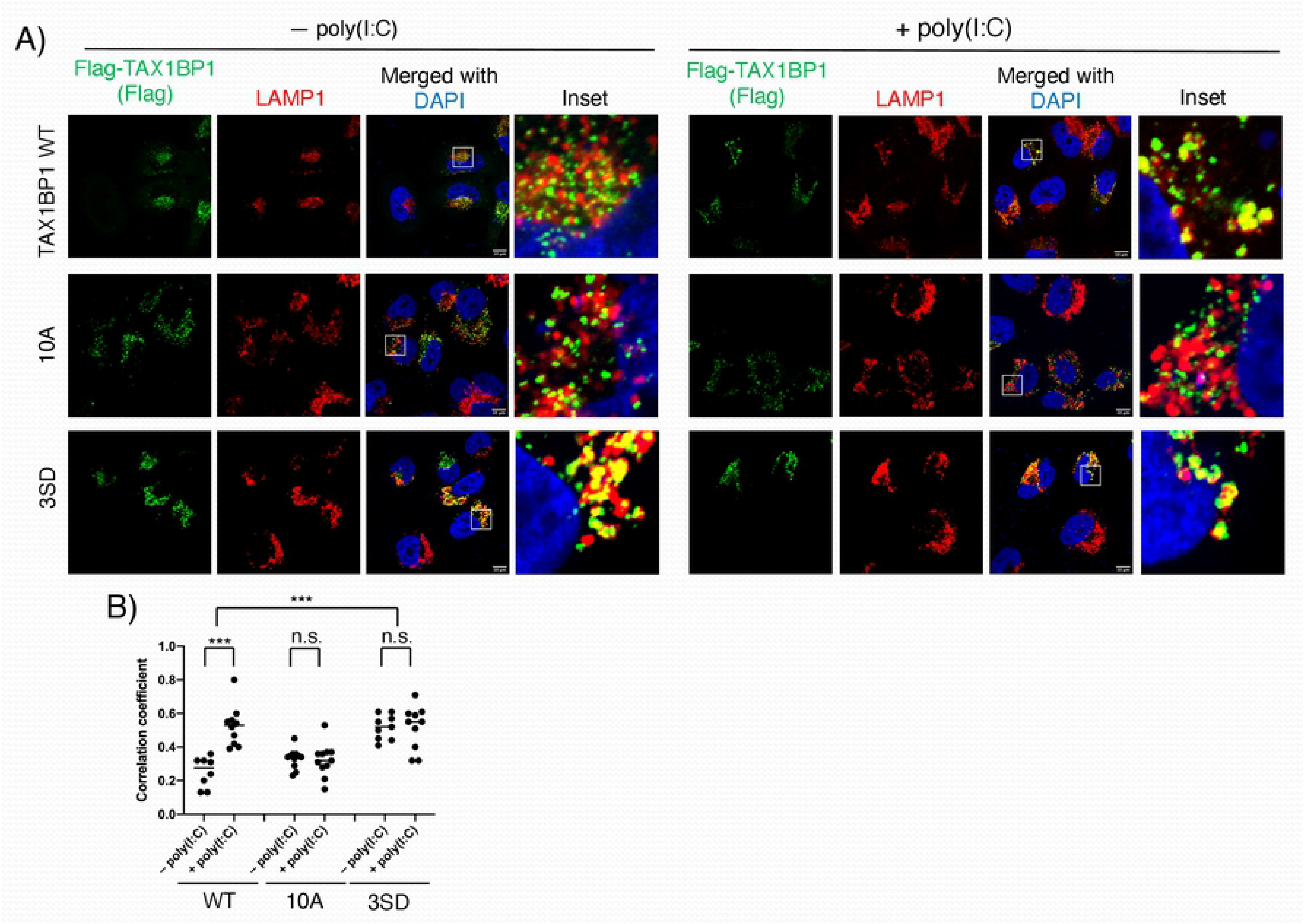
Phosphorylation of TAX1BP1 is required for localization to autolysosomes. (A) Immunofluorescence assays. *TAX1BP1* KO HeLa cells were transfected with Flag-TAX1BP1 WT, 10A or 3SD (S254D, S593D and S666D) and 24 h later transfected with 2.5 μg/ml poly(I:C) for 6 h in the presence of 20 μM leupeptin. Scale bar, 10 μm. (B) Pearson’s correlation coefficient was calculated to measure co-localization between TAX1BP1 and LAMP1 in 8-12 cells randomly selected from each sample. Unpaired Student’s *t*-test, \*\*\**P* <0.001, n.s.=not significant.

### TAX1BP1 clears MAVS aggregates in a phosphorylation-dependent manner

Thus far our experiments have established that TAX1BP1 undergoes autophagic degradation during RNA virus infection, and this process is dependent on its phosphorylation. It remains unclear what cargo are recruited to autophagosomes by TAX1BP1 for the homeostatic control of the RLR pathway. We previously demonstrated that TAX1BP1 interacts with the mitochondrial adaptor MAVS, and targets MAVS for degradation [29]. Basal expression of MAVS protein is elevated in *TAX1BP1* KO cells and the half-life of MAVS is significantly increased in the absence of TAX1BP1 [29]. Dysregulated MAVS expression in *TAX1BP1* KO cells results in increased type I IFN and apoptosis in response to RNA virus infection [29]. Upon activation of RLR signaling MAVS forms large aggregates with properties of amyloid fibers and prions including: 1) formation of fiber-like polymers, 2) ability to “infect” the endogenous protein and convert it to aggregate forms, 3) resistance to protease digestion and 4) resistance to detergent solubilization [43]. A common approach to analyze MAVS aggregates is SDD-AGE (semi-denaturing detergent agarose gel electrophoresis), which can detect large polymers between 200-4000 kDa [43, 44]. Therefore, SDD-AGE was utilized to analyze MAVS aggregates in WT and *Taxlbp1*^−/−^ MEFs infected with Sendai virus (SeV). In WT MEFs, SeV infection triggered the formation of MAVS aggregates as expected (Fig 7A). Remarkably, MAVS aggregates were spontaneously produced in uninfected *Taxlbp1*^−/−^ MEFs, at levels greater that WT MEFs infected with SeV (Fig 7A). SeV infection further increased MAVS aggregates in KO cells (Fig 7A). We next asked if overexpression of TAX1BP1 could clear MAVS aggregates, and if phosphorylation of TAX1BP1 played a role in this function. Overexpression of MAVS yielded aggregates as detected by SDD-AGE; however, the phosphomimetic 3SD TAX1BP1 mutant but not the phosphorylation mutant 10A inhibited MAVS aggregates (Fig 7B). Furthermore, the phosphomimetic 3SD TAX1BP1 mutant was more effective than WT TAX1BP1 in suppressing MAVS-induced IFN activation (Fig 7C). However, the TAX1BP1 phosphorylation mutant 10A was impaired in the inhibition of MAVS-IFN induction (Fig 7C).

**Fig. 7.**
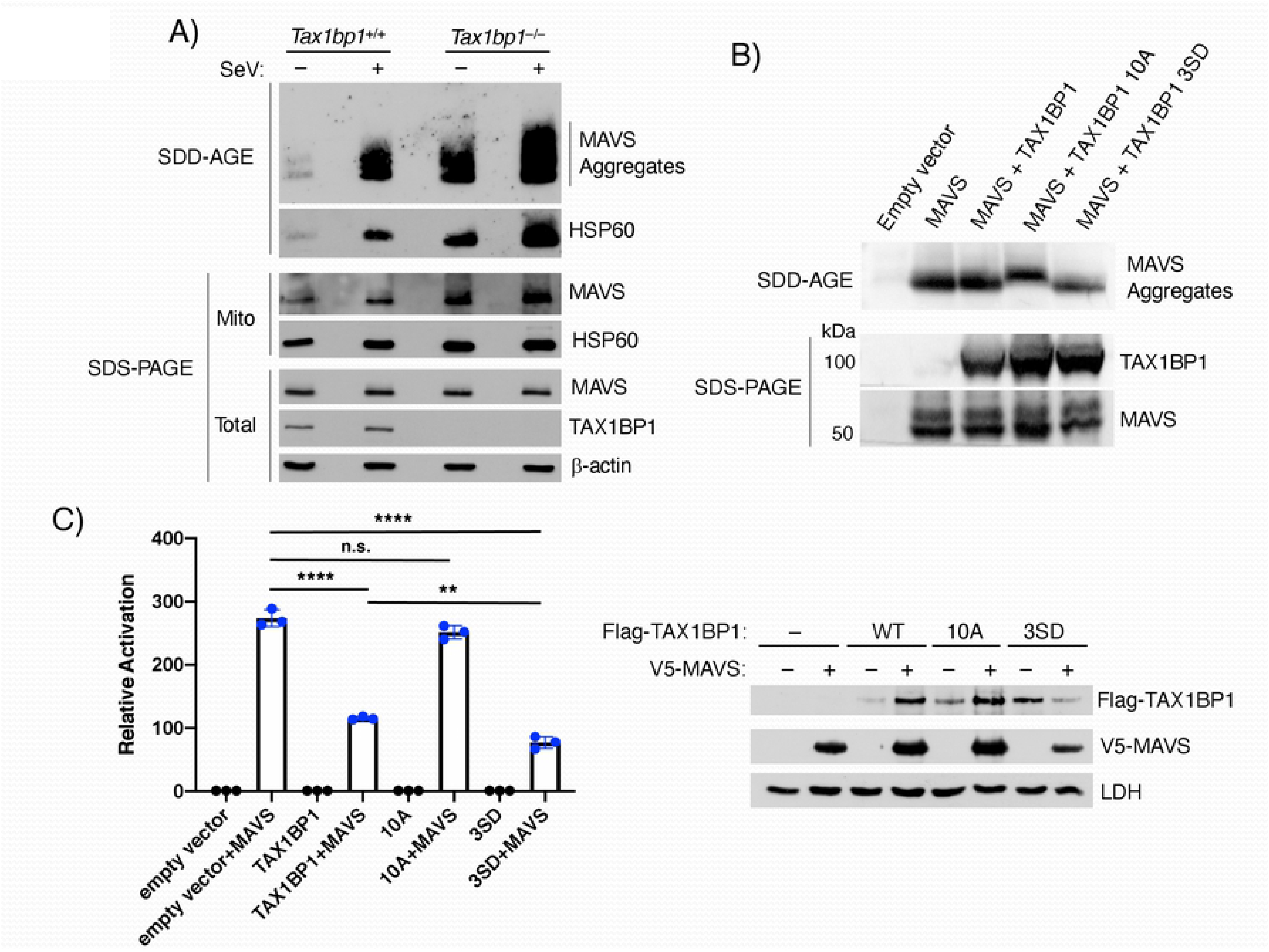
TAX1BP1 promotes MAVS degradation via aggrephagy. (A) Semi-denaturing detergent agarose gel electrophoresis (SDD-AGE) analysis of MAVS protein. Crude mitochondria were isolated from *Tax1bp1^+/+^* and *Tax1bp1*^−/−^ MEFs infected with Sendai Virus (SeV) (25 HA units/ml) for 6 h, and extracts separated on SDD-AGE and SDS-PAGE gels and immunoblotted with the indicated antibodies. (B) 293 T cells were transfected with the indicated plasmids and lysates subjected to SDD-AGE for MAVS aggregates and SDS-PAGE to examine expression of TAX1BP1 and MAVS. (C) Dual luciferase reporter assays. 293T cells were co-transfected with V5-MAVS and Flag-TAX1BP1 WT, 10A or 3SD at a ratio of 1:8 along with interferon β (IFNβ) promoter-driven firefly luciferase and thymidine kinase (TK) promoter-dependent *Renilla* luciferase reporter plasmids for 24 h. The data are presented as mean ± standard deviation of biological triplicates. The remaining cell lysates were subjected to immunoblotting with anti-Flag, anti-V5 and anti-LDH antibodies. One-way ANOVA with Dunnett’s post hoc test, \*\*\*\**P* <0.0001, n.s.=not significant. Unpaired Student’s *t*-test, **, *p* < 0.01.

## Discussion

In this manuscript we have found that the selective autophagy receptor TAX1BP1 can be phosphorylated by noncanonical and canonical IκB kinases, with 13 putative IKKi-induced phosphorylation sites in TAX1BP1 identified by mass spectrometry analysis. TAX1BP1 phosphorylation triggers its autophagosomal degradation with serines 254, 593 and 666 playing the most critical roles. Whereas IKKi and TBK1 regulate basal TAX1BP1 phosphorylation and degradation, IKKα is required for RNA virus-mediated TAX1BP1 autophagosomal degradation. Furthermore, the core autophagy factor ATG9 plays critical roles in both basal and virus-triggered TAX1BP1 degradation. Finally, we found that TAX1BP1 serves as a phosphorylation-dependent aggrephagy receptor for MAVS, and TAX1BP1-deficient cells exhibit a spontaneous accumulation of MAVS aggregates.

Phosphorylation plays important functional roles in the regulation of selective autophagy receptors. TBK1 phosphorylates OPTN, NDP52 and p62/SQSTM1 during bacterial infection and mitophagy to enhance Ub binding and cargo recruitment to autophagosomes [45–50]. Although TBK1 was shown to interact with TAX1BP1 in a previous study [51], it was not examined whether TBK1 played any role in TAX1BP1 autophagy function. Our data indicate that TBK1 and IKKi can phosphorylate TAX1BP1, but surprisingly both were dispensable for RNA virus-induced TAX1BP1 degradation (Fig 3). Instead, IKKα was required for TAX1BP1 degradation by RNA virus infection (Fig 4). In this regard, we previously found that IKKα interacts with TAX1BP1 and phosphorylates TAX1BP1 on serines 593 and 666 for the inhibition of NF-κB signaling [39], however the effect of this phosphorylation on TAX1BP1 autophagy function was not examined in our previous study. TBK1 and IKKi share identical phosphorylation motifs which overlap with IKKα and IKKβ substrate specificities [52–54]. IKK phosphorylation sites usually have acidic or phosphorylated amino acids both amino-terminal and carboxyl-terminal to the phosphorylation site. Our bioinformatics and experimental approaches have identified S254 and S593 in TAX1BP1 as phosphorylation sites for IKKi and IKKα (Fig 1) [39]. S666 does not conform to an IKK consensus site and thus may be phosphorylated by a kinase other than IKKi or IKKα. S254 and S593 are positioned immediately downstream of coiled-coil domains 1 and 3 respectively, which overlap with self-oligomerization regions of TAX1BP1 [23, 55]. However, TAX1BP1 phosphorylation at S254 or S593 did not enhance its dimerization or binding to LC3 or MAVS (Figs S5-S7). Rather, IKKα and IKKi-mediated TAX1BP1 phosphorylation promotes its lysosomal localization (Figs 6 and S8), possibly due to conformational changes in TAX1BP1 that may facilitate its trafficking and/or binding to other factors needed for autophagosome/autolysosome targeting.

TAX1BP1 degradation during RNA virus infection or poly(I:C) transfection is inhibited by Baf A1 treatment or by genetic deletion of the core autophagy factor ATG9 (Fig 5), suggesting that TAX1BP1 degradation is mainly mediated by the classical autophagy pathway. However, we noticed there was not a complete block in virus-induced TAX1BP1 degradation in *Alg3*^−/−^ MEFs (Fig 5B). These data indicate that a fraction of TAX1BP1 can still be degraded during virus infection in the absence of the LC3 lipidation machinery. Two recent studies have described lysosomal degradation pathways involving TAX1BP1 that are independent of ATG7 and LC3 lipidation [42, 56]. Future studies should determine if TAX1BP1 is also degraded by these LC3-independent lysosomal targeting pathways during RNA virus infection.

MAVS forms large prion-like aggregates during RNA virus infection that propagate downstream signaling for the activation of TBK1 and IRF3, and type I IFN. However, it remains poorly understood how MAVS aggregates are resolved once viral infections are cleared to suppress inflammation and autoimmunity. Indeed, MAVS aggregates can be detected in peripheral blood mononuclear cells (PBMCs) of systemic lupus erythematosus (SLE) patients and are associated with increased levels of type I IFN [57]. The E3 ligase MARCH5 inhibits RNA virus-induced MAVS aggregates by ubiquitinating MAVS to trigger its proteasomal degradation [58]. The deubiquitinase YOD1 also antagonizes MAVS aggregation by cleaving K63-linked poly Ub chains on MAVS [59]. MAVS is also negatively regulated by autophagic degradation although the mechanisms remain poorly understood. The E3 ligases RNF34 and MARCH8 have been shown to ubiquitinate MAVS to promote NDP52-dependent autophagic degradation of MAVS [19, 20]. Our results indicate that TAX1BP1 serves a nonredundant role in preventing the spontaneous formation of MAVS aggregates and also inhibits MAVS aggregates formed during RNA virus infection (Fig 7). Therefore, we conclude that TAX1BP1 functions as an aggrephagy receptor for MAVS, which is congruent with a recent study that described TAX1BP1 as an aggrephagy receptor in the brain [38]. It remains to be determined if TAX1BP1 aggrephagy function extends to other innate immune signaling pathways in addition to the RLR pathway.

In summary, we describe a novel regulatory role for phosphorylation in the regulation of TAX1BP1 autophagosomal degradation and aggrephagy function. Since several viral proteins (e.g., HTLV-1 Tax [55], human papillomavirus E2 [60] and measles virus nucleoprotein [61]) interact with TAX1BP1, it will be interesting in future studies to examine if viruses exploit TAX1BP1 phosphorylation to inhibit MAVS and RLR signaling.

## Materials and Methods

### Cell culture

Human embryonic kidney 293T (HEK293T) and DLD-1 cells were purchased from the American Type Culture Collection (ATCC). *TAX1BP1* knockout (KO) HeLa cells were provided by Dr. Richard Youle [36]. *Tax1bp1*^−/−^ MEFs were described previously [24]. *Ikkα*^−/−^, *Ikkβ*^−/−^ and *Ikkp*^−/−^ MEFs were provided by Dr. Michael Karin and described previously [39]. *Atg3*^−/−^ [62], *Ikki*^−/−^ [63] and *Ikki^−/−^Tbk1^−/−^* [64] MEFs were obtained from the indicated sources. Cell lines and MEFs were cultured in Dulbecco’s Modified Eagle Medium (DMEM) supplemented with 10% fetal bovine serum, streptomycin and penicillin at 37°C and 5% CO_2_. The cell lines were tested for mycoplasma contamination using MycoAlert^®^ Mycoplasma Detection kit (R&D Systems) and if necessary cultured in the presence of Plasmocin™ treatment or Plasmocin™ prophylactic (InvivoGen). Transient transfection with plasmids was performed using GenJet version II (SignaGen Laboratories), and transfection with poly(I:C) was performed using Lipofectamine 2000 reagent (Invitrogen) following the manufacturer’s instructions.

### Virus infections

Cells were starved for 1 h in serum-free DMEM and inoculated with VSV-GFP or SeV for 1 h at the indicated multiplicity of infection (MOI) in serum-free DMEM, and further incubated in complete DMEM for the indicated times.

### Immunological assays

Antibodies used in immunological assays including immunoblotting, immunoprecipitation (IP), and indirect immunofluorescence (IFA) are listed in Supplemental Table 1. Cells were lysed in RIPA buffer (50 mM Tris [pH 7.4], 150 mM NaCl, 1% Igepal CA-630, and 0.25% deoxycholate) containing a protease inhibitor cocktail and protein phosphatase inhibitors, including 10 mM NaF and 5 mM Na_3_VO_4_. For immunoblotting, cell lysates were separated on SDS-PAGE, transferred to nitrocellulose or polyvinylidene difluoride membranes, and immunoblotted with appropriate primary antibodies diluted in SuperBlock™ (PBS) blocking buffer (Thermo Fisher Scientific). Following incubation with horseradish peroxidase-labeled appropriate secondary antibody, immunoreactive bands were visualized by an enhanced chemiluminescence (ECL) reagent on an X-ray film. ImageJ software (NIH) was used to quantify the intensities of bands. For IP, total cell extracts were incubated with Flag (L5) or V5-antibody-conjugated beads overnight. Immunoprecipitants were washed with RIPA buffer, followed by elution of bound proteins with 1x SDS sample buffer. For IFA, cells grown on a coverslip (and transfected) were fixed in Image-iT™ fixative (Thermo Fisher Scientific) and permeabilized in 0.5% Triton X-100 prepared in PBS. Following incubation with SuperBlock™ PBS blocking buffer for 1 h at room temperature, coverslips were incubated with primary antibodies, washed with PBS, and then incubated with appropriate fluorescent dye-conjugated secondary antibodies. Coverslips were mounted in ProLong™ Gold Antifade Mounting medium (Thermo Fisher Scientific) containing 4’, 6-diamidino-2-phenylindole (DAPI) on glass slides and cells were imaged on a Zeiss 700 confocal laser scanning microscope with a 63x oil-immersion objective and Zen software. Pearson’s correlation coefficient was calculated to measure co-localization between TAX1BP1 and LAMP1 using Coloc2 (ImageJ).

### Nucleic acid manipulation

All polymerase chain reaction (PCR) amplification and site-directed mutagenesis were performed using Platinum™ *Pfx* or SuperFi™ DNA polymerase (Thermo Fisher Scientific). Subcloning of open reading frames (ORFs) and their derivatives into expression plasmids was conducted using appropriate restriction enzyme sites (Supplemental Table 2). Small guide RNAs (gRNAs) for each gene were selected using Deskgen software, synthesized by Integrated DNA Technologies, and cloned into plentiCRISPR v2-puro [65] or plentiCRISPR v2-blasticidin (a gift from Mohan Babu, Addgene plasmid #83480) using BsmBI. Oligonucleotides are listed in Supplemental Table 3.

### CRISPR/Cas9-mediated gene knockout

CRISPR/Cas9-mediated genetic ablation of the indicated genes in DLD-1 cells was performed as previously described [66]. Antibiotic-resistant individual clones were isolated by limiting dilution and genomic DNA purified and subjected to Sanger DNA sequencing. TBK1 and IKKi gRNAs cloned in pLentiCRISPRv2 were kindly provided by Dr. Fangfang Zhou [67].

### *In vitro* kinase and phosphatase assays

Purified recombinant human TAX1BP1 protein (catalog # P01; Abnova) was incubated with purified GST-tagged IKKi (catalog # PV4875; Thermo Fisher Scientific) in buffer A containing 20 mM Tris [pH 7.5], 10 mM MgCl_2_, 1 mM EGTA, 1 mM Na_3_VO_4_, 5 mM β-glycerophosphate, 2 mM DTT, 0.02% Triton X-100 and 200 μM ATP for 10 min at 30°C or with purified hexahistidine-tagged TBK1 (catalog # PV3504; Thermo Fisher Scientific) in buffer B containing 50 mM Tris [pH 7.5], 10 mM MgCl_2_, 2 mM DTT, 0.025% Triton X-100 and 200 μM ATP for 20 min at 30°C. The reaction was terminated by boiling in 1x SDS-sample buffer for 5 min. For dephosphorylation of TAX1BP1, lysates from 293T transfected cells were incubated with 800U λ-phosphatase at 30°C for 30 min and then subjected to SDS-PAGE analysis.

### Mass spectrometry (MS) analysis

Coomassie stained gel pieces were de-stained and subjected to reduction (5 mM DTT for 45 min at 60°C) and alkylation (20 mM iodoacetamide for 20 min at room temperature in the dark). Samples were subsequently proteolyzed with 10 ng trypsin (Promega)/μl overnight at 37°C. Dry extracted peptides after clean-up were re-suspended in 8 μl 0.1% formic acid. Titanium dioxide was used for phosphopeptide enrichment. Protein identification by liquid chromatography tandem mass spectrometry (LC-MS/MS) analysis of peptides was performed using an LTQ Orbitrap Velos MS (Thermo Fisher Scientific) interfaced with a nanoAcquity LC system (Waters, Corp.). Peptides were fractionated by reverse-phase HPLC on a 75 μm x 15 cm PicoFrit column with a 15 μm emitter (New Objective) in-house packed with Magic C18AQ (Michrom BioResources, Inc.) using 0-60% acetonitrile/0.1% formic acid gradient over 70 min at 300 nl/min. Eluting peptides were sprayed directly into an LTQ Orbitrap Velos at 2.0 kV. Survey scans were acquired from 350-1,800 m/z with up to 10 peptide masses individually isolated with a 1.9 Da window and fragmented (MS/MS) using a collision energy of 40 and 30s dynamic exclusion. Precursor and the fragment ions were analyzed at 30,000 and 7500 resolution, respectively. Peptide sequences were identified from isotopically resolved masses in MS and MS/MS spectra extracted with and without deconvolution using Thermo Scientific MS2 processor and Xtract software. Data was searched for in the human RefSeq database, with oxidation on methionine (variable), deamidation NQ (variable), phosphoSTY (variable) and carbamidomethyl on cysteine as (fixed) modifications, using Proteome Discoverer 1.3 software.

### Semi-denaturing detergent agarose gel electrophoresis (SDD-AGE)

SDD-AGE was performed as previously performed [68]. Briefly, crude mitochondria isolated by differential centrifugation were resuspended in 1x sample buffer (0.5x Tris-borate-EDTA [TBE], 10% glycerol, 2% SDS, and 0.0025% bromophenol blue) and loaded onto a 1.5% agarose gel. After electrophoresis in the running buffer (1x TBE and 0.1% SDS) for 1 h with a constant voltage of 100 V at 4 °C, proteins were transferred to a nitrocellulose membrane for immunoblotting. SDD-AGE with transfected MAVS and TAX1BP1 was performed similarly but with whole cell lysates.

### Live-cell imaging

A total of 5×10^4^ WT or *Atg3*^−/−^ MEFs were infected with VSV-GFP (MOI=0.1) and live-cell imaging was performed with an Incucyte S3 Live-Cell Analysis System (Essen BioScience). Images were acquired every two hours using phase contrast and green channels with a 10x objective in triplicate. GFP fluorescence was quantified as green object count per image normalized to phase area confluence.

### Dual luciferase reporter assays

Cells were transfected with the desired plasmids together with IFNβ-Luc and the *Renilla* reporter pTK-RLuc as an internal control. After 24 h, cells were lysed in passive lysis buffer (Promega) and subjected to dual-luciferase assays as recommended by the manufacturer (Promega). Luminescence was measured using a GloMax luminometer (Promega). Results are presented as the relative firefly luciferase activity over the *Renilla* luciferase activity.

### NanoBiT assays

NanoBiT assays were performed as described by the manufacturer (Promega). Briefly, 293T cells were seeded in a 6-well plate and transfected with pairs of NanoBiT constructs. After 24 h, cells were collected, washed two times in PBS [pH 7.4], and resuspended in 1 ml of Opti-MEM I Reduced Serum Media (Thermo Fisher Scientific). 100 μl of cell suspension was transferred to a white-walled 96-well plate in triplicate, and 20 μl of Nano-Glo^®^ Luciferase substrate furimazine (Promega) diluted in PBS at a ratio of 1:100 was added to each well. After incubation for 5 min at room temperature, luminescence was measured by a GloMax luminometer (Promega).

### Reagents

Poly(I:C), Bafilomycin A1, IKK inhibitor VII, and leupeptin were purchased from MilliporeSigma. λ-phosphatase was from New England Biolabs. Calyculin A was from Cell Signaling. SeV was purchased from Charles River Laboratories.

### Statistical analysis

Data are presented as mean ± standard deviation from a representative experiment with triplicate samples. Statistical analysis was performed in GraphPad Prism 8 and indicated in the Figure legends.

## Acknowledgements

We thank Drs. Hong-Gang Wang and Victor Jin for *Atg3*^−/−^ MEFs, Dr. Tom Maniatis for *Ikki*^−/−^ MEFs, Drs. Thomas Decker and Shizuo Akira for *Ikki^−/−^Tbk1^−/−^* MEFs, Dr. Michael Karin for *Ikkα*^−/−^, *Ikkβ^−/−^ and Ikkγ*^−/−^ MEFs, Dr. Richard Youle for *TAX1BP1* KO HeLa cells, Dr. Fangfang Zhou for TBK1 and IKKi CRISPR/Cas9 plasmids and Dr. Siddharth Balachandran for VSV-GFP.

## Figure legends

**Fig. S1 A kinase-dead IKKi mutant is impaired in TAX1BP1 phosphorylation.**

293T cells were transfected with Flag-TAX1BP1 and either Flag-IKKi or Flag-IKKi K38A. Lysates were subjected to immunoblotting with anti-Flag antibody.

**Fig. S2 Mapping of TAX1BP1 sites phosphorylated by IKKi.**

293T cells co-transfected with Flag-TAX1BP1 WT and variants (10A and its variants, which were rescued at the indicated residues) together with Flag-IKKi for 24 h, and the cell extracts were immunoblotted with anti-Flag antibody.

**Fig. S3 IKKi and TBK1 are not involved in virus infection-induced TAX1BP1 degradation.**

Two different IKKi and TBK1 double KO (dKO) DLD-1 cell lines were infected with VSV-GFP for 24 h at an MOI of 0.1, and cell extracts were immunoblotted with anti-TAX1BP1, p62 and LDH antibodies.

**Fig. S4 TAX1BP1 can interact with members of the mammalian autophagy-related gene 8 (mATG8) family.**

Co-immunoprecipitation (IP) assays. 293T cells were co-transfected with each of the V5-tagged mATG8 members, including MAP1LC3A (LC3A), MAP1LC3B (LC3B), MAP1LC3C (LC3C), GABARAP, GABARAPL1 (GEC1), and GABARAPL2 (GATE16), together with Flag-TAX1BP1 and 24 h later lysed, and the lysates were subjected to IP using anti-Flag antibody-conjugated agarose (L5 beads, BioLegend). The IP complex and lysates were separated on SDS-PAGE and immunoblotted with anti-Flag or V5 antibody.

**Fig. S5 Phosphorylation does not regulate mATG8 binding of TAX1BP1.**

Co-immunoprecipitation (IP) assays. 293T cells were co-transfected with V5-LC3B or V5-GEC1 together with Flag-TAX1BP1 WT and variants and 24 h later lysed, and the lysates were subjected to IP using anti-Flag antibody-conjugated agarose (L5 beads, BioLegend). The IP complex and lysates were separated on SDS-PAGE and immunoblotted with anti-Flag or V5 antibody.

**Fig. S6 TAX1BP1 binding to MAVS does not require its phosphorylation.**

Co-immunoprecipitation (IP) assays. 293T cells were co-transfected with V5-tagged MAVS together with Flag-TAX1BP1 (WT and indicated variants) and 24 h later lysed, and the lysates were subjected to IP using anti-V5 antibody-conjugated agarose. The IP complex and lysates were separated on SDS-PAGE and immunoblotted with anti-Flag or V5 antibody.

**Fig. S7 Phosphorylation does not affect TAX1BP1 dimerization**.

NanoBiT-based protein fragment complementation assays (PCA). NanoBiT assays were performed as previously described [68]. (A) 293T cells were co-transfected with Nano luciferase Large BiT (LgB)-fused TAX1BP1 together with Small BiT (SmB)-fused HaloTag (HT) or TAX1BP1 and 24 h later transfected with or without 2.5 μg/ml poly(I:C) for an additional 6 h. The luminescence was measured using GloMax (Promega) after adding furimazine (Promega), a cell-permeable substrate of Nano luciferase. RLU was calculated by normalizing the data by the value of the pair of HT-SmB and LgB-TAX1BP1 (no poly(I:C) treatment). The data presented are mean ± the standard deviation (SD) of biological triplicates. Unpaired Student’s *t*-test, ***, *p* < 0.001. (B) 293T cells were co-transfected with pairs of LgB and SmB-fused TAX1BP1 (WT, 10A and 3SD) proteins and 24 h later measured for their luminescence as above. In addition, the TAX1BP1 mutant lacking amino acids 321-420, which are essential for TAX1BP1 dimerization, was included as a control. The data presented are mean ± SD of experimental triplicates.

**Fig. S8 Overexpression of IKKα and IKKi promotes TAX1BP1 localization to autolysosomes.** *TAX1BP1* KO HeLa cells were co-transfected with Flag-TAX1BP1 WT along with empty vector, IKKα, IKKβ or IKKi, and 24 h later immunostained with Flag and LAMP1 antibodies. Leupeptin (20 μM) was added to the cultures to prevent autophagic degradation of TAX1BP1. A representative confocal image of each sample is presented. Scale bar, 10 μm.

